# Protein Kinase A activity is regulated by actomyosin contractility during cell migration and is required for durotaxis

**DOI:** 10.1101/392399

**Authors:** Andrew J. McKenzie, Tamara F. Williams, Kathryn V. Svec, Alan K. Howe

## Abstract

Dynamic subcellular regulation of Protein kinase A (PKA) activity is important for the motile behavior of many cell types, yet the mechanisms governing PKA activity during cell migration remain largely unknown. The motility of SKOV-3 epithelial ovarian cancer (EOC) cells has been shown to be dependent on both localized PKA activity and, more recently, on mechanical reciprocity between cellular tension and extracellular matrix (ECM) rigidity. Here, we investigated the possibility that PKA is regulated by mechanical signaling during migration. We find that localized PKA activity in migrating cells rapidly decreases upon inhibition of actomyosin contractility (specifically, of myosin ATPase, ROCK (Rho kinase), or MLCK (myosin light chain kinase) activity). Moreover, PKA activity is spatially and temporally correlated with cellular traction forces in migrating cells. Additionally, PKA is rapidly and locally activated by mechanical stretch in an actomyosin contractility-dependent manner. Finally, inhibition of PKA activity inhibits mechanically-guided migration, also known as durotaxis. These observations establish PKA as a locally-regulated effector of cellular mechanotransduction and as a regulator of mechanically-guided cell migration.

## Introduction

Cells sense, respond, and contribute to the mechanical properties of the extracellular matrix (ECM) by exerting actomyosin-dependent contractile force on integrin-based adhesive contacts and sensing counter-tension through mechano-chemical systems (1–7). Integrin engagement and clustering initiates the generation of cellular forces through actomyosin contractility (8–10), which in turn promotes maturation and strengthening of adhesive contacts, thereby providing counter-tension to the force of protrusive actin polymerization within leading edge lamellipodia and large-scale contractility to pull the cell body in the direction of migration (11–18). The distribution of contractile forces within a migrating cell generates subcellular areas with varying degrees of intracellular tension, countered by both internal cytoskeletal scaffolds and by matrix adhesions (7, 19). Intracellular contractile forces regulate myriad aspects of cell migration (4, 13, 20–22) and, during the migration of many cell types, traction force tends to be highest within the leading edge, often within the lamella just behind actively protruding lamellipodia (23–27).

The cAMP-dependent protein kinase (PKA) is an important regulator of myriad targets involved in cell migration and cytoskeletal dynamics (28, 29) and localized activation of PKA signaling, facilitated by A-kinase anchoring proteins (AKAPs), is necessary for the motile behavior of numerous cell types (30–39). However, the mechanisms controlling PKA activation during migration remain unclear.

Previously, we showed that efficient migration of SKOV-3 human ovarian cancer cells requires localized PKA activity within the leading edge (32). More recently, we demonstrated that SKOV-3 cell migration is also governed by the mechanical microenvironment; specifically, that cell contractility and migration positively correlate with ECM stiffness and that directional increases in ECM tension promote SKOV-3 cell durotaxis (27). In the present work, we explored the possibility that localized PKA activity in migrating cells might be regulated by mechanical signaling and cell-matrix tension.

## Results

### PKA is activated at the leading edge during cell migration but not at the periphery during cell spreading

In an attempt to facilitate the investigation of the regulation of PKA within the leading edge of migrating cells, we performed live-cell FRET microscopy using the PKA biosensor pmAKAR3 to examine the PKA activity in the periphery of live, spreading SKOV-3 cells shortly after plating onto fibronectin-coated surfaces, as this periphery is often used as a model for the migratory leading edge (*e.g*. (40)). However, little to no significant PKA activity was observed in spreading cells (Fig. 1A & C; Supplemental Movie S1), even though robust, dynamic PKA activity was seen within the leading edge of migrating cells (Fig. 1B & C; Supplemental Movie S1), consistent with prior reports (32). This was somewhat surprising, given that prior investigations have reported, using biochemical methods, some degree of activation of PKA early upon integrin engagement and during cell spreading (41, 42). It is important to point out, here, that the pmAKAR3 biosensor used throughout this study (and many others (31–33, 35)) is targeted to the plasma membrane *via* a C-terminal CAAX box (derived from K-Ras; (43)), and thus only reports membrane-proximal events. Thus, there may be other pools of PKA that are activated during spreading that cannot directly affect and alter this membrane-bound biosensor. In any event, these data suggest that spreading cells are not a suitable experimental system for studying regulation of PKA within the leading edge.

**Figure 1.**
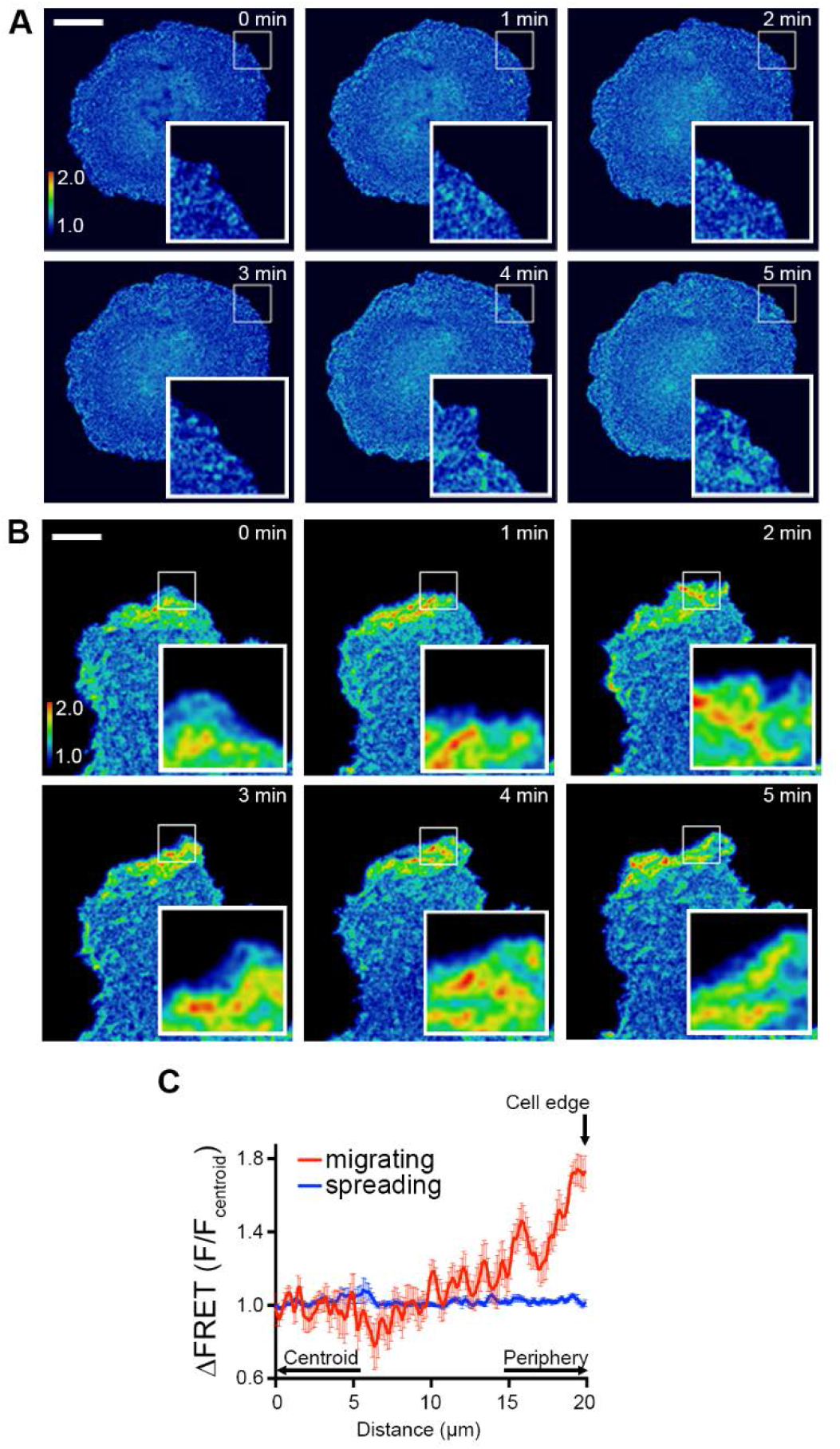
PKA activity is increased at the leading edge of migrating cells but not at the periphery of spreading cells. SKOV-3 cells expressing pmAKAR3 were plated on fibronectin-coated glass coverslips 10 min (A) or 4 h (B) after plating, then imaged *via* FRET microscopy. Representative pseudocolored FRET/CFP images of spreading (A) and migrating (B) cells are shown and areas of protrusion are magnified in the insets (bar = 10 μm). (C) Linescan analysis of the change centroid-to-front FRET ratio was performed on migrating (red) and spreading (blue) cells values are displayed as mean ± SEM. (*n* = 14 cells for each condition). Images were acquired every 60 sec.

### Integrin engagement is not sufficient to elicit peripheral/leading edge PKA activity

Over the course of additional experimentation and optimization of conditions for visualizing localized PKA activity during migration, we noticed that activation of PKA in the leading edge was common to many modes of migration under a variety of culture conditions, *e.g*. signaling events were seen in the presence or absence of serum, before and after serum starvation, and with and without supplemental growth factor, albeit with differences in signaling event morphology and kinetics (Supplemental Movie S2). This suggested that, rather than being instigated by soluble cues from culture media, the mechanisms governing localized PKA activity during migration might be governed by something intrinsic to cells as they move.

One process common to most if not all modes of migration is the engagement of integrins by the ECM and the subsequent formation of adhesive complexes such as focal contacts and focal adhesions (44). Importantly, engagement of integrins has also been implicated in activation of PKA (28, 30, 33, 41, 42, 45–47). We therefore investigated whether there were any spatio-temporal links between leading edge PKA signaling events and the formation of focal adhesion. For this, we imaged cells co-expressing pmAKAR3 and mCherry-paxillin, to visualize PKA activity and focal adhesion dynamics, respectively. Paxillin was used as a marker of focal adhesions because it recruits to focal adhesions early in the maturation process and remains through the entirety of the adhesion lifetime (48). Visual assessment of numerous images suggested no close or obvious spatial correlation between leading edge PKA events and paxillin-containing focal adhesions (Fig. 2A). To further assess any spatiotemporal correlation between focal adhesion dynamics and PKA activity, leading edge focal adhesions were tracked over time and an interrogation region of interest (ROI) 50% larger than the area of the adhesion was used to track pmAKAR3 FRET ratios under the same cellular regions (Fig. 2B). Paxillin displayed full adhesion lifecycles including assembly, peak, and disassembly phases (Fig. 2C, red line) while PKA activity showed little, if any, variation in the same regions. Additionally, ROIs were placed around areas of dynamic leading edge PKA activity events and used to measure corresponding mCherry-paxillin intensities (Fig. 2D). Interestingly, there was no apparent co-variation between the intensity of mCherry-paxillin and peak PKA activity events (Fig. 2E). Similar results were seen using mCherry-FAK as distinct focal adhesion marker (Supplemental Fig. S1). To ensure that an increase in focal adhesion intensity correlated with the recruitment of signaling events known to be regulated by focal adhesion dynamics, cells were co-transfected with mCherry-paxillin and YFP-dSH2 ((49) yellow fluorescent protein fused to two tandem Src SH2 domains that bind phospho-tyrosine. We saw strong co-variations of paxillin intensity and the intensity of YFP-dSH2 during the assembly, peak, and disassembly of focal adhesions (Fig. 2F). Quantification of Pearson’s correlation coefficients showed a strong positive correlation between mCherry-paxillin and YFP-dSH2, but no correlation between pmAKAR3 FRET ratios and mCherry-paxillin (Fig. 2G). While we cannot rule out a role for small and/or transient focal complexes that are below the threshold of our detection, these results demonstrate that leading edge PKA dynamics are not spatiotemporally correlated with the onset, maturation, or dissolution of mature focal adhesions in migrating cells. This suggests that leading edge PKA activity is regulated through a mechanism dependent on, but downstream of and spatially removed from, integrin-mediated focal adhesion formation.

**Figure 2.**
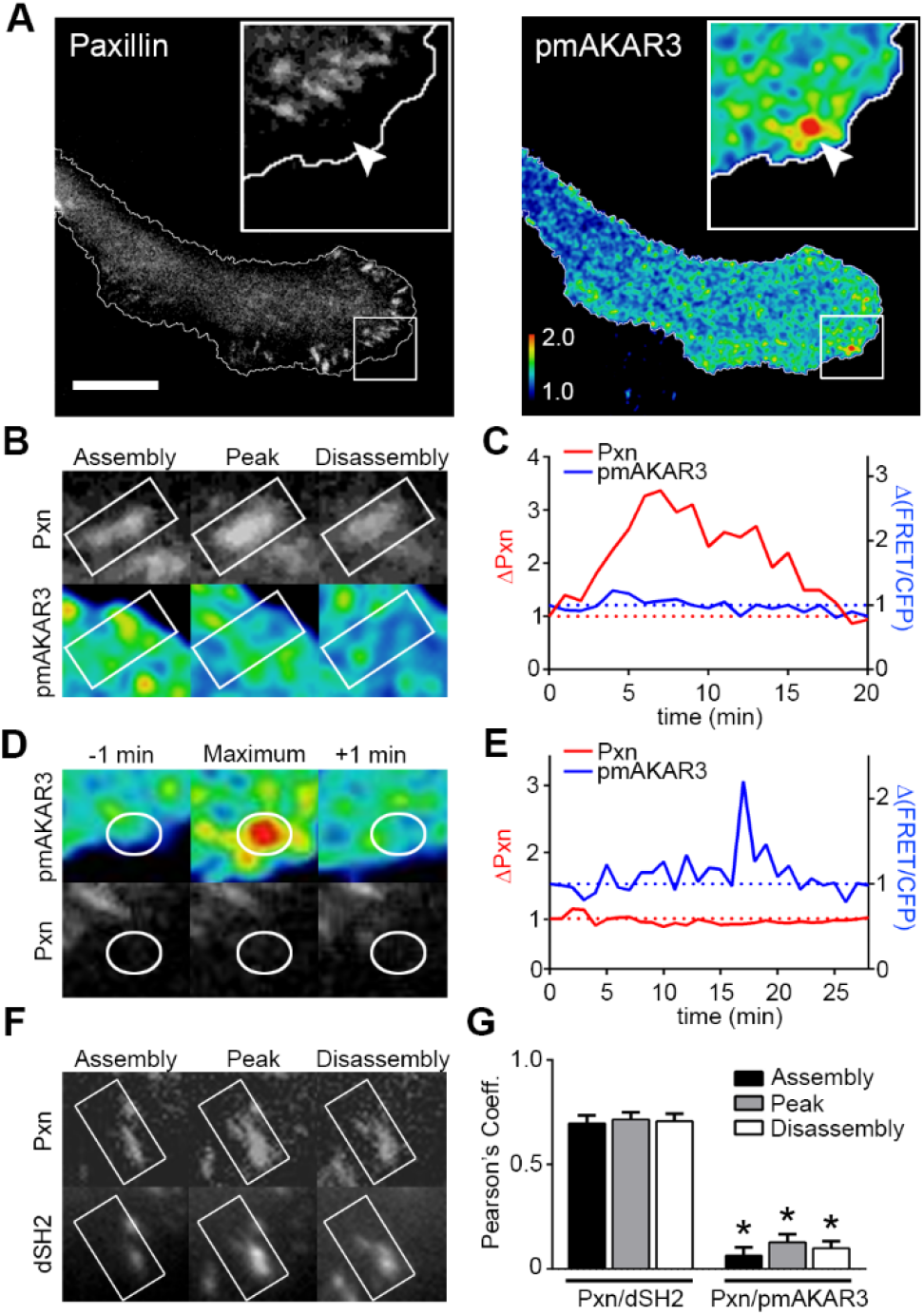
Leading edge PKA activity does not correlate with focal adhesion dynamics. (A) Migrating SKOV-3 cells co-expressing pmAKAR3 and mCherry-paxillin were plated on fibronectin coated imaging dishes and representative near-simultaneous paxillin (Pxn) and pseudocolored FRET/CFP (pmAKAR3) images from live-cell microscopy experiments are shown. Arrow indicates location of peak FRET/CFP signal and corresponding location on mCherry-paxillin image. The leading edge is magnified in the insets are shown with cell outlines plotted for reference (bar = 10 μm). (B) Images of a single paxillin-containing focal adhesion during assembly, peak, and disassembly (top) with overlapping pseudocolored FRET/CFP (bottom) are shown. (C) The changes in either paxillin fluorescence intensity or FRET ratios were plotted over time. (D) Images of dynamic PKA activity and the corresponding paxillin images are shown with ROI plotted as reference. (E) The changes in either FRET ratios or paxillin fluorescence intensity were plotted over time. (F) YFP-dSH2 and paxillin intensities were tracked over time during focal adhesion assembly, peak, and disassembly are shown. (G) Pearson’s coefficients of the covariance of paxillin with either YFP-dSH2 (Pxn/dSH2) or pmAKAR3 (Pxn/pmAKAR3) are during focal adhesion assembly, peak and disassembly are shown as mean ± SEM (Pxn/dSH2, *n* = 10; Pxn/pmAKAR3, *n* = 13; * = p<0.001 for each phase of focal adhesion lifetime).

### Leading edge PKA activity is regulated by actomyosin contractility

In further consideration of what might regulate PKA activity during migration, we began to consider a cellular characteristic that is dependent on integrin-mediated adhesion, intrinsic to many cells across many modes of migration, and not typically associated with cell spreading – namely, actomyosin-dependent cellular contractility (1, 13, 17, 50, 51). Intracellular contractile forces regulate diverse aspects of signaling and cytoskeletal dynamics during cell migration (4, 13, 20–22) and, importantly, these forces tend to be highest within the leading edge (23–27), placing them in the correct subcellular location to affect PKA activity.

To investigate whether cellular contractility affected PKA dynamics, we imaged PKA activity in live cells before and after addition of blebbistatin, an inhibitor of the myosin II-ATPase. Upon addition of blebbistatin, localized and dynamic activity of PKA in the leading edge rapidly and significantly diminished (Fig. 3A and B; Supplemental Movie S3). Similar effects were seen upon treatment of migrating cells with inhibitors of Rho-kinase (H-1152 or Fasudil/HA-1077; Fig. 3C & Supplemental Movie S4) or myosin light chain kinase (ML-7; Supplemental Movie S4), both of which contribute to the phosphorylation and activation of the myosin light chain (MLC) of myosin II to promote contractility (52, 53).

**Figure 3.**
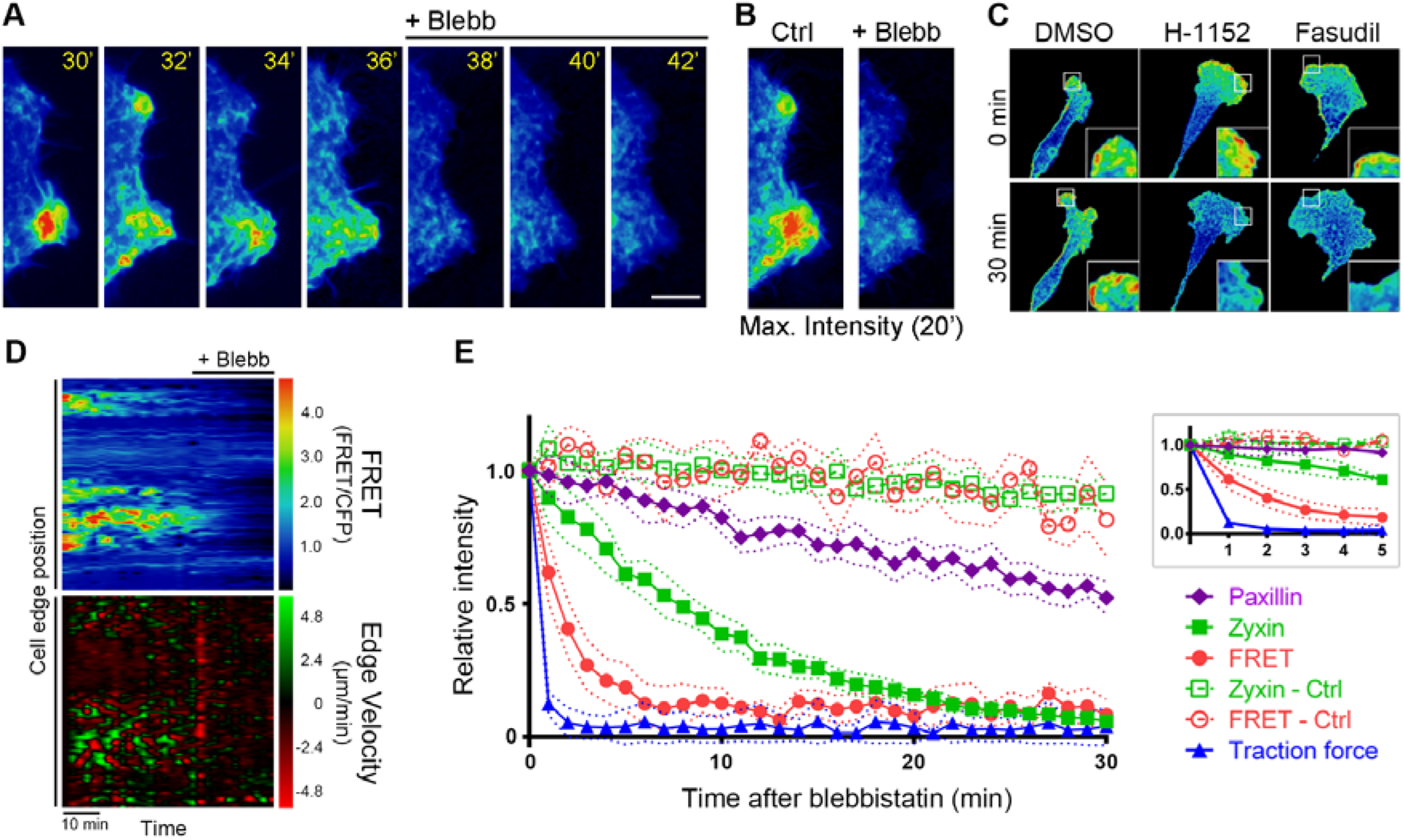
Actomyosin contractility regulates leading edge velocity and PKA activity. (A) A migrating SKOV-3 cell expressing pmAKAR3 was plated on fibronectin-coated a glass-bottom imaging dish and monitored by live-cell microscopy before and after treatment with 25 μM blebbistatin (*Blebb*). Representative pseudocolored FRET/CFP images (corresponding to the boxed region in Supplemental Movie S3) are shown. (B) A maximum intensity projection of cumulative PKA activity in ten images (2 min apart) before and after treatment with blebbistatin. (C) Representative images of pmAKAR3-expressing SKOV-3 cells migrating on fibronectin-coated dishes before (*0 min*) and 30 min after treatment with 0.1 % vol/vol DMSO, 1 μM H-1152, or 10 μM fasudil. The leading edge is magnified in the insets (scale bar = 10 μm). (D) QuimP11 software (54) was used to generate maps of PKA activity (top) and edge-velocity (bottom) within a 10 μm band along the leading edge before and after treatment with blebbistatin (*Blebb*). (E) SKOV-3 cells expressing mCherry-paxillin, mCherry-zyxin, or pmAKAR3 (*FRET*) migrating on fibronectin-coated glass dishes, or cells plated on 125 kPa fluorescent nanosphere-functionalized hydrogels (*Traction force*) were treated with 25 μM blebbistatin (or with 0.1% vol/vol DMSO; *Ctrl*) at time=0. Capturing images every 60 sec, the fluorescence intensity of paxillin or zyxin within focal adhesions, PKA activity (*via* pmAKAR FRET signal), or traction force was measured, normalized to values at t = 0, and plotted. The graph depicts mean values ± SEM (*n*_Paxillin-Blebb_ = 211 adhesions from 7 cells; *n*_Zyxin-Blebb_ = 156 adhesions from 5 cells; *n*_FRET_ = 30 linescans from 6 cells; *n*_Zyxin-Ctrl_ = 125 adhesions from 5 cells; *n*_FRET-Ctrl_ = 35 linescans from 7 cells; *n*_Traction Force_ = average traction from 6 cells). The inset shows the first five minutes of the time course at higher resolution; these data were used to calculate one-phase half-life (t½) or apparent t½ decay values for each signal using GraphPad Prism.

Inhibition of actomyosin contractility is associated with discrete morphological effects, including cessation of leading edge dynamics and eventual dissolution of focal adhesions, so it is possible that the loss of PKA is coupled to one of those events. To better assess the kinetics of loss of PKA activity upon inhibition of contractility, we generated morphodynamic maps of protrusion/retraction velocities and PKA activity along the leading edge using QuimP11 edge tracking and sampling software (54). As demonstrated previously (33), edge velocity and leading edge PKA activity are spatiotemporally correlated in actively migrating cells (Fig. 3D). Importantly, we also observed a rapid, coincident decrease in edge dynamics and leading edge PKA activity upon treatment with blebbistatin (Fig. 3D). While the observations in Fig. 2 suggest that there is no spatial correlation between PKA signaling events and focal adhesion dynamics, focal adhesions are important centers of signal transduction and their dissolution upon inhibition of contractility would be expected to disrupt that signaling. Specifically, if PKA activity was dependent on signaling from intact focal adhesions, we would predict that disassembly of focal adhesions would precede loss of PKA activity. Thus, we assessed the kinetics of inhibition of PKA activity relative to focal adhesion disassembly by measuring the rate of blebbistatin-induced loss of fluorescent signal intensity of two focal adhesion markers: zyxin, a mechano-sensitive protein that leaves focal adhesion rapidly upon loss of contractility, and paxillin, which leaves ‘relaxed’ focal adhesions much more slowly and with apparent zero-order kinetics (17, 48, 55). As expected, paxillin intensity within focal adhesions decreased slowly but steadily after blebbistatin treatment (Fig. 3E), with an apparent half-life of 52.69 ± 3.35 min, while zyxin intensity decreased rapidly and exponentially, with a half-life of 7.63 ± 0.66 min (Fig. 3E; Supplemental Movie S5), consistent with prior observations of the higher dependence of zyxin on mechanical forces for residence within focal adhesions (17, 55–58). Interestingly, PKA activity decreased even more rapidly after blebbistatin treatment than focal zyxin intensity (Fig. 3E), with a half-life of ~50 sec (0.83 ± 0.09 min). These observations are consistent with our observation of the lack of spatiotemporal correlation between focal adhesion dynamics and peripheral PKA signaling events and suggest a closer relative coupling of PKA to the contractile state of the cell. With this in mind, we then determined the kinetics of loss of cellular traction force after blebbistatin treatment using traction force microscopy on cells adhered to fibronectin-coated polyacrylamide hydrogels functionalized with fluorescent nanospheres (27). As expected, cellular traction force decreased very rapidly after addition of blebbistatin, with a t½ of 0.37 ± 0.05 min, (Fig. 3E & Supplemental Movie S6). While this t½value is likely to be erroneously high, given that the rate of image acquisition for these experiments (1 frame/min) results in an image interval that far exceeds the apparent t½, it is consistent with published reports (*i.e*. (17, 55, 58)). More importantly, this decrease was far more rapid than the loss of paxillin or even zyxin from focal adhesions but only slightly more rapidly than loss of PKA activity. Collectively, these data demonstrate that, during cell migration, regulation of PKA activity within the leading edge is kinetically coupled to and dependent on actomyosin contractility.

### Spatial distribution of cellular traction forces and PKA activity during cell migration

If leading edge PKA is regulated by actomyosin contractility, then one might expect PKA activity and cellular contractile forces to be coincident in migrating cells. To investigate this correlation directly, SKOV-3 cells expressing pmAKAR3 and migrating on nanosphere-functionalized hydrogels were analyzed by simultaneous FRET and traction force microscopy (TFM) in the absence and presence of blebbistatin (Fig. 4). In the absence of blebbistatin, migrating cells exhibited high traction forces at sites of protrusion which overlapped with areas of high PKA activity (Fig. 4A) and linescan analysis through cellular leading edges confirmed that the radial increase in PKA activity from cell center to periphery overlapped with peripherally-increasing traction forces (Fig. 4B). Of note, traction forces were significantly lower in spreading cells compared to traction forces in migrating cells (Supplemental Fig. S3), consistent with the aforementioned lack of peripheral PKA activity in spreading cells (Fig. 1). Importantly, both cellular traction forces and PKA activity decreased dramatically upon treatment with blebbistatin (Fig. 4A) and the residual pockets of contractility were no longer spatially correlated with residual PKA activity (Fig. 4C).

**Figure 4.**
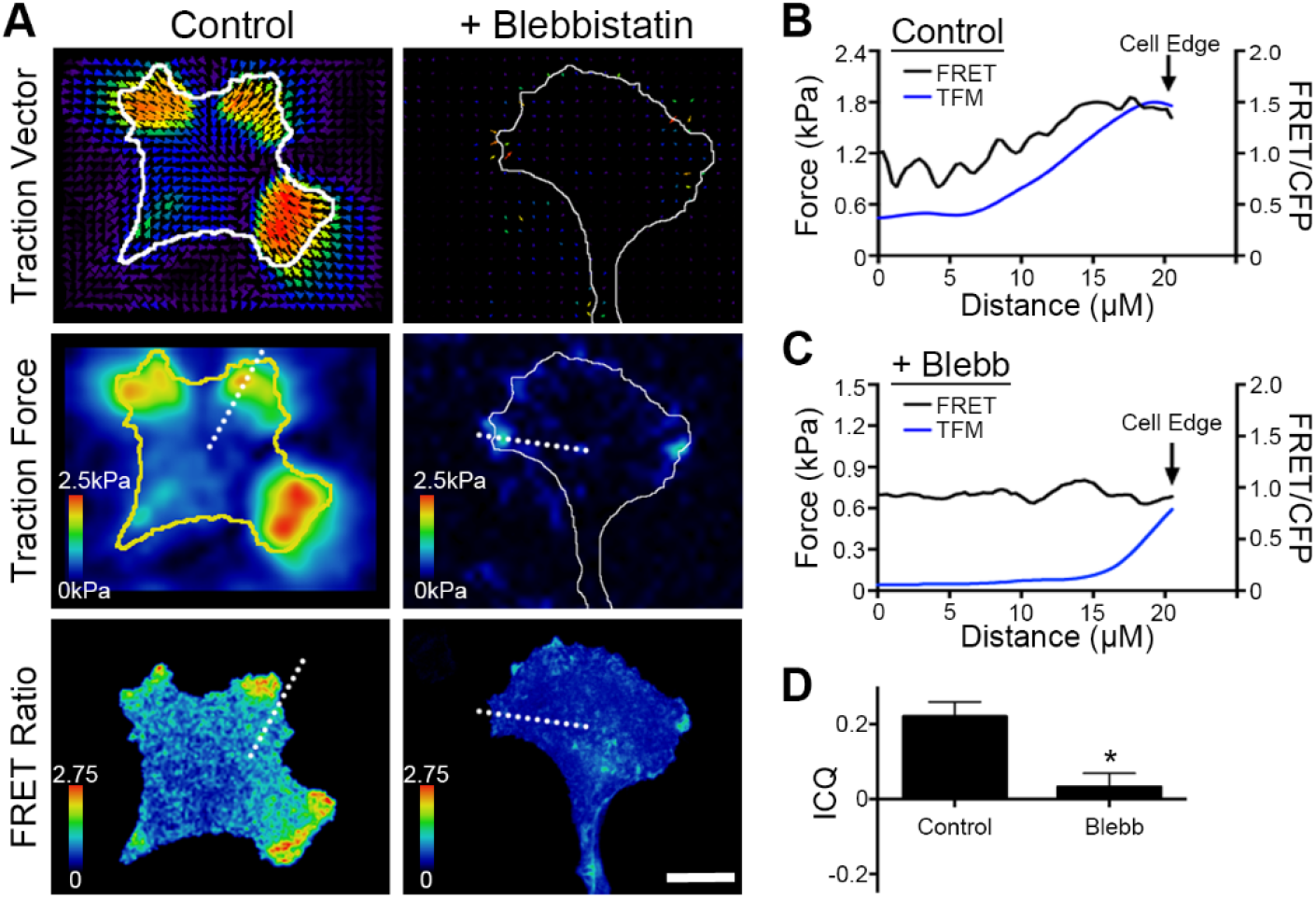
PKA activity and cellular traction forces are spatiotemporally correlated. (A) Cells expressing pmAKAR3 were plated on fibronectin-coated polyacrylamide hydrogels surface-conjugated with fluorescent 0.2 μm nanospheres and imaged by FRET microscopy and traction force microscopy after 20 min treatment with DMSO (*Control*) or 25 μM blebbistatin. The top panels show bead displacement/traction vector maps and the middle panels show traction force maps (with cell outlines plotted for reference) while the bottom panels show time-matched FRET/CFP images (bar = 10 μm). (B, C) Linescan analyses (dashed white lines in panel A) of both traction forces and FRET intensity from the cell centroid to the cell periphery for cells either control (B) or blebbistatin-treated (C) cells. (D) Pixel-by-pixel image correlation analysis was performed using intensity correlation analysis software (see *Materials and Methods*) to generate an intensity correlation quotient (ICQ). ICQs between traction force maps and PKA activity maps are summarized as mean ± SEM (*n* = 7 cells for each condition; * = fv To formally quantify the extent *p*<0.001).

To formally quantify the extent to which PKA activity and traction forces overlap in migrating cells, TFM and FRET images were subject to co-localization and intensity correlation analysis using intensity correlation quotients (ICQ). ICQ provide a single value indicating the covariance of two signals that can be used for statistical comparison (59, 60). Mean ICQ values of −0.05 to +0.05 indicate random distribution of the two signals; values less than −0.05 indicate mutual exclusion; values between +0.05 and +0.1 indicate moderate covariance; and values > 0.1 indicate strong covariance. Under control conditions, the mean ICQ for traction forces and PKA activity was 0.226 ± 0.022 and was significantly reduced to 0.032 ± 0.021 when the cells were treated with blebbistatin (mean ± SEM, Fig. 4D). These data demonstrate that PKA activity and traction forces are spatially coincident in migrating cells. Coupled with the earlier observations that inhibition of actomyosin contractility significantly inhibits leading edge PKA activity, these observations strongly suggest that PKA activity is locally regulated by a contractility-dependent mechanotransduction pathway during cell migration.

### PKA is activated by mechanical stretch in an actomyosin-dependent manner

Given the demonstrated requirement of cellular tension for localized activation of PKA during migration, we wondered whether PKA might be locally activated by acute increases in cellular tension. To test this, cells expressing pmAKAR3 alone or pmAKAR3 along with either mCherry or mCherry fused to the PKA-inhibitor protein (mCh-PKI; (32)) were plated on fibronectin-coated hydrogels to promote cell migration. Then, the hydrogel under individual cells was stretched (perpendicular to the axis of cell migration) with a microneedle to elicit a rapid, local increase in matrix rigidity and cell-matrix tension (Fig. 5A). Upon application of directional stretch to control cells, there was a rapid, robust, and localized increase in PKA activity in the direction of stretch (Fig. 5B-D). Importantly, this acute mechanical activation of PKA was completely inhibited in cells co-expressing mCh-PKI (Fig. 5C & E) or treated with blebbistatin prior to stretch (Fig. 5E). These results show that acute mechanical stimulation can activate PKA activity in a manner that requires actomyosin contractility.

**Figure 5.**
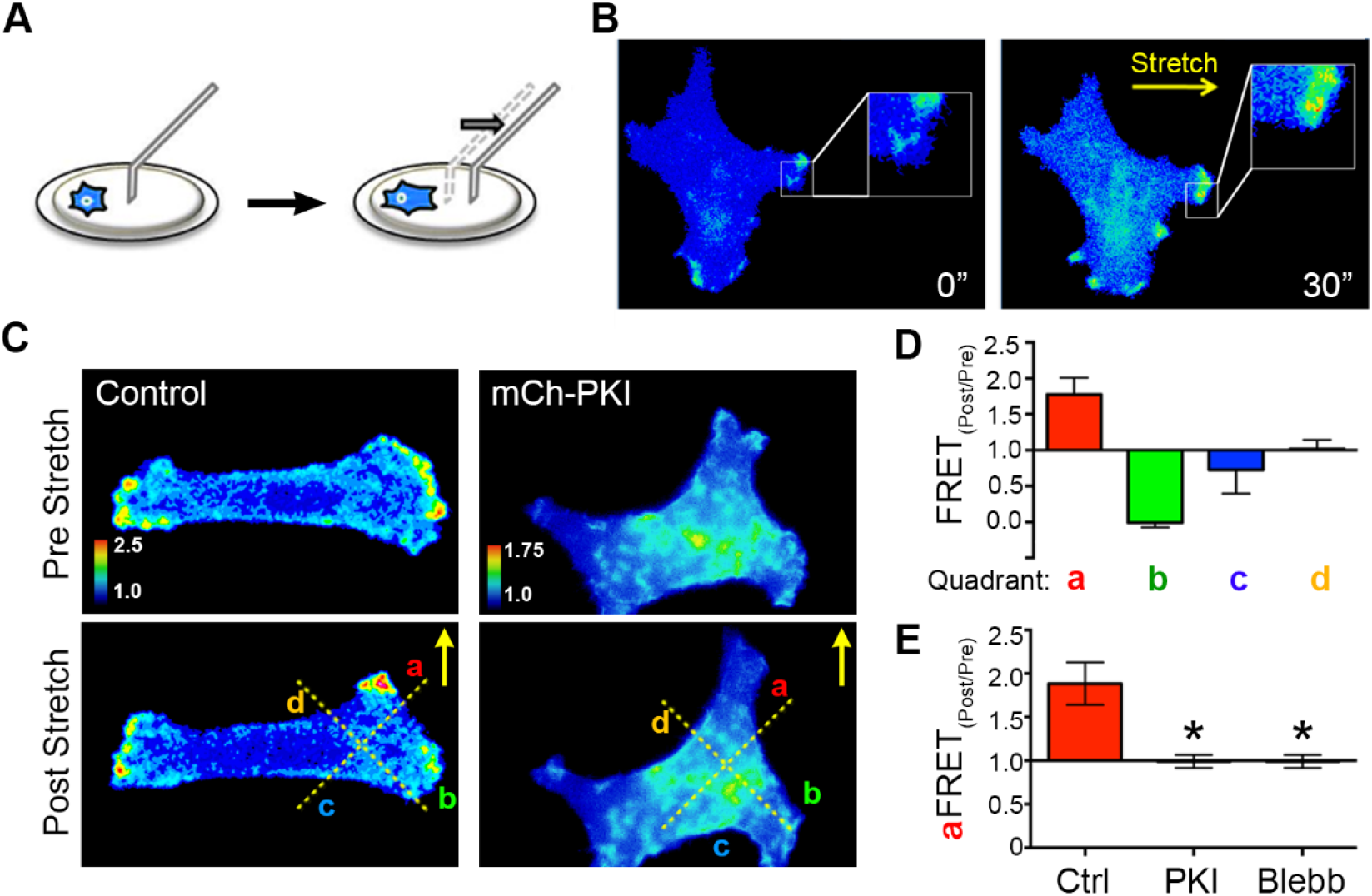
PKA is activated upon acute mechanical stretch and is required for SKOV-3 cell durotaxis. (A) Schematic of the technique used to impart acute, directional tension on individual cells using a glass microprobe to deform the underlying hydrogel (see text for details). (B) An SKOV-3 cell expressing pmAKAR3 was plated on a fibronectin-coated hydrogel for 4 h, then imaged by FRET microscopy before and 30 sec after application of mechanical stretch in the indicated direction. (C) SKOV-3 cells co-expressing pmAKAR3 and either mCherry (Control) or mCherry-PKI (mCh-PKI) were cultured and stimulated by directional stretch (indicated by the arrows) as in panel (B). PKA activity (from pmAKAR3 FRET) was imaged before and ~30 sec after stretch. Images were sectioned into quadrants (*a* through *d*), depicted by dotted lines, for further analysis. (D) The change in PKA activity before and after stretch (FRET_Post/Pre_) of controls cells was calculated in quadrants proximal (quadrant *a*), orthogonal (quadrants *b,d*) or contralateral (quadrant *c*) to the stretch (mean ± SEM; *n* = 6) (E) The change in PKA activity was monitored in quadrant *a* (proximal to stretch) in SKOV-3 cells either co-expressing pmAKAR3 and mCh-PKI or mCherry as a control, or cells expressing pmAKAR3 and pre-treated with 25 μM blebbistatin (Blebb) for 10 min (mean ± SEM; *n* = 5 cells *per* condition; * = *p*< 0.05 (one-way *ANOVA*)).

### PKA activity is required for durotaxis in SKOV-3 cells

Previously, we have shown that efficient migration in SKOV-3 cells is dependent upon PKA activity (32). More recently, we showed that the migration of these cells is strongly influenced by the mechanical microenvironment and that these cells exhibit durotaxis, or mechanically-guided migration towards regions of increased ECM rigidity (27). Given the current observations connecting cellular tension & stretch to activation of PKA, we investigated whether the durotactic migration of SKOV-3 might be similarly dependent on PKA activity. To this end, control cells or cells expressing mCh-PKI were cultured to migrate on fibronectin-coated hydrogels, subject to directional stretch as described above, and monitored for durotactic response (see *Materials and Methods*). While controls cells exhibited robust durotaxis in response to acute, directional stretch, inhibition of PKA activity dramatically decreased durotactic efficiency (Fig. 6A-C; Supplemental Movie S7). Taken together, these observations demonstrate that PKA activity is mechanically regulated during cell migration, is activated upon acute, mechanical cell stretch, and is required for durotaxis.

**Figure 6.**
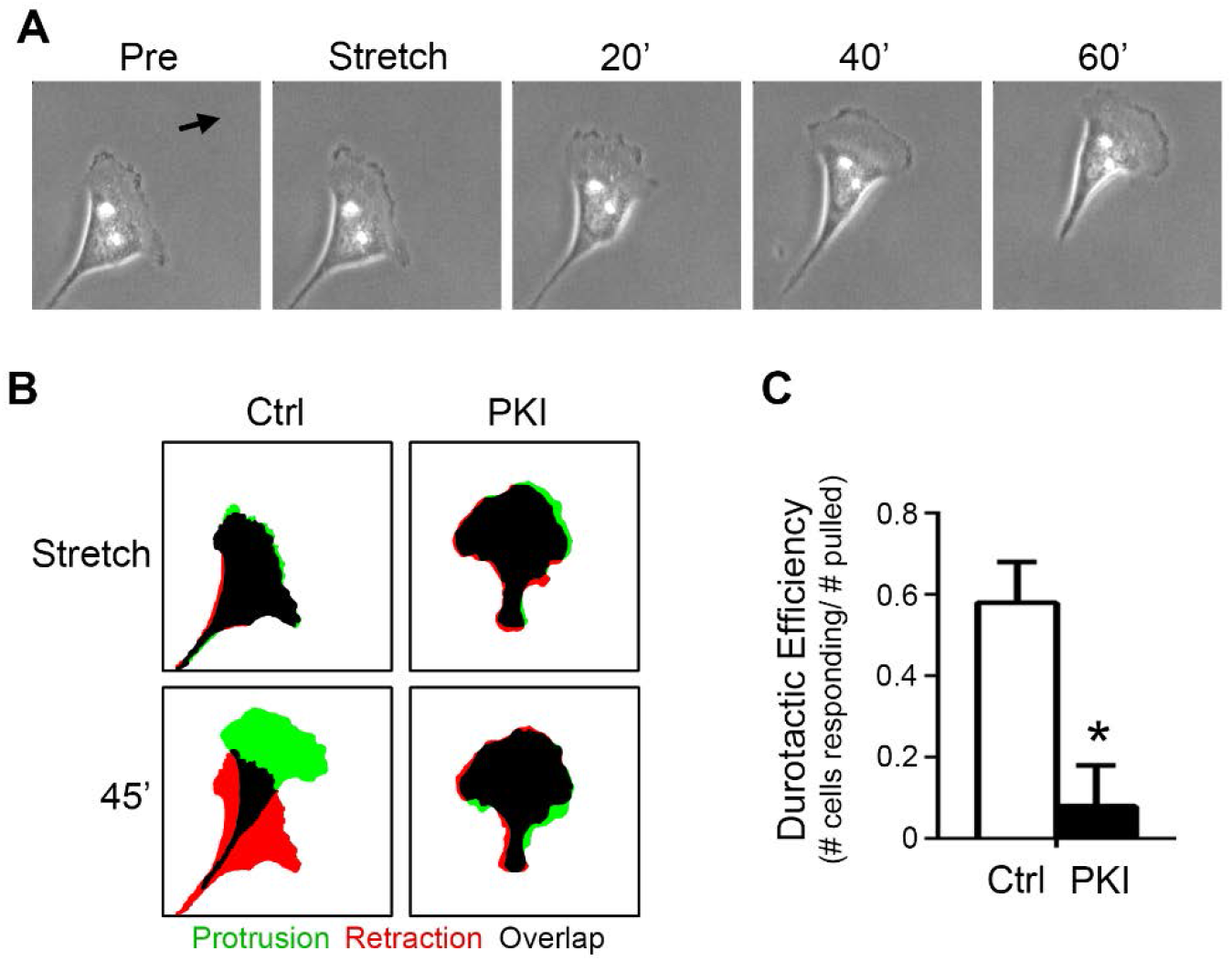
PKA is required for SKOV-3 cell durotaxis. (A) An SKOV-3 cell plated on a 25 kPa, fibronectin-coated hydrogel for 4 h was monitored by live-cell phase-contrast microscopy before and after durotactic stimulation (in the direction indicated by the arrow); shown are images captured one minute before (*Pre*) application of stretch, one minute after application (*Stretch*), and every 20 min thereafter are shown. (B) Control cells or cells expressing mCherry-PKI (*PKI*) were cultured and stimulated to invoke durotaxis as described for panel (A). Protrusion/retraction analysis maps (PRAMs) were generated from manually thresholded and outlined phase-contrast images taken 1 min before and 1 min after stretch (*Stretch*) and 1 min and 45 min after stretch (45’) to identify regions of protrusion, retraction, and overlap (green, red, and black respectively). (C) Durotactic efficiency (see *Materials & Methods* for details) was calculated for control and mCherry-PKI-expressing cells (*n* = 8 for each condition; * = *p*<0.01).

## Discussion

The mechanosensitivity of cAMP/PKA signaling is well-supported by the literature. For example, the cAMP cascade was the first mechanosensitive signaling cascade ever to be described (61). Also, the levels of cAMP and PKA activity have been shown to vary as a function of the mechanical tension of fibroblasts embedded in collagen gels (62). Furthermore, direct application of mechanical force through integrin-mediated adhesive contacts rapidly activates cAMP and PKA signaling and PKA-dependent transcription (45, 46, 63). Finally, mechanical signals from fluid flow activate PKA in an ECM-specific manner (64). While these and other studies firmly establish PKA as a mechano-responsive target, it is important to note that they all assessed global rather than subcellular activation of cAMP/PKA and did not follow PKA dynamics in migrating cells.

Here, we demonstrate a link between cellular tension and regulation of subcellular PKA activity during cell migration. We show that leading edge PKA activity correlates with the spatial distribution of cellular traction forces and that disruption of actomyosin contractility uncouples the spatial correlation between cellular forces and PKA activity. We establish that leading edge PKA activity is regulated by cellular tension and show that locally-applied mechanical forces elicit localized increases in PKA activity. Furthermore, we demonstrate PKA activity is required for the durotactic response to directional mechanical cues, thereby expanding this enzyme’s well-established role as a regulator of other modes of cell migration (30–33, 35). An intriguing recent report studying cells migrating from open to confined two-dimensional spaces showed that PKA activity was down-regulated during this transition in a manner dependent on Piezo 1-mediated Ca^2+^ influx and, importantly, that leading edge PKA activity appeared to increase upon treatment of cells with blebbistatin (65). These latter observations are in direct contrast to the results reported herein, which demonstrate a decrease in PKA activity upon treatment with not only blebbistatin, but also with other inhibitors of actomyosin contractility (Y-27632, H-1152, fasudil/HA1077, and ML7; Fig. 3). The reasons for this disparity are currently unknown but may include differences in cell lines and culture conditions. The prior study saw increased PKA activity after blebbistatin treatment (in both confined and unconfined cells) using CHO-K1 cells and a derivative line expressing the α4 integrin (CHO-α4WT), adhered to surfaces coated with 20 μg/ml fibronectin. These are notable, yet modest differences compared to the current work, which uses SKOV-3 cell adhered to fibronectin at 10 μg/ml. While it will be important to keep these differences in mind for future studies, it is perhaps more important to appreciate that, together, the two reports firmly establish PKA as a target for mechanical regulation during migration.

While previous work has shown activation of leading edge PKA activity to be integrin-mediated (34, 35, 41, 42, 47, 64, 66, 67), our current observations establish that this activity is both spatially and temporally distinct from sites of integrin-dependent focal adhesion assembly. It is important to reiterate that we cannot rule out a contribution of direct, integrin-mediated signaling events arising from focal complexes that are below the level of detection using the current methods. However, we contend that the circuitry for localized activation of PKA is spatially, temporally, and thus biochemically separate from mature focal adhesions. Moreover, we do *not* contend that focal adhesions are wholly unnecessary for regulating PKA during migration. As the nexus between the actin cytoskeleton and matrix-bound integrins, focal adhesions are principal mediators of mechanical signaling. However, the relative kinetics of PKA inactivation and focal adhesion disassembly suggest that PKA is regulated by a tension-dependent aspect/characteristic of intact focal adhesions (e.g. a mechanically regulated enzymatic activity or protein-protein interaction) rather than bulk focal adhesion assembly or disassembly.

Though the current work establishes the connection between leading edge PKA activity and actomyosin contractility, the exact molecular mechanism coupling cellular tension to localized PKA activity remains to be elucidated. Given that PKA has been shown to physically anchor directly to integrin cytoplasmic tails (34), it is possible that mechanical tension on integrins may be transmitted directly to PKA and somehow activate it, perhaps by pulling the regulatory subunits apart from the catalytic subunits. While intriguing, Occam’s razor would suggest that this is an unlikely mechanism. A more likely hypothesis is that canonical activators of the cAMP/PKA pathway (G-protein coupled receptors, adenylyl cyclases, or phosphodiesterases) are locally and mechanically regulated during cell migration. Importantly, while GPCRs are well-established mediators of mechanotransduction (68), the vast majority of stretch-sensitive GPCRs couple to Gαq heterotrimers and would require a more circuitous route to activate PKA. However, there are reports of stretch-sensitive GPCRs that couple to Gαs (68) and experiments imparting tension across fibronectin-coated magnetic beads demonstrated that force application to integrins led to activation PKA in a Gαs-dependent manner (45, 63). Those reports, along with the current observations, give rise to the intriguing hypothesis that adhesive and/or cytoskeletal elements may exhibit mechanically-controlled cross-talk, directly or indirectly, with GPCRs – a possibility with important consequences for cell migration and myriad other cellular functions.

## Materials and methods

### Reagents and cell culture

Human epithelial ovarian cancer (SKOV-3) cells were purchased from American Type Culture Collection, authenticated at the UVM Advanced Genomic Technology Core, and were maintained in a humidified incubator at 37°C containing 5% CO_2_ in DMEM supplemented with 10% fetal bovine serum. Cells were trypsinized and split 1:5 every 3–4 days to avoid reaching confluence. Cells were transfected using Fugene6 (Promega) according to the manufacturer’s protocol. In brief, cells were plated into 35 mm dishes at 60–70% confluence the day prior to transfection so that they were 75-85% confluent at the time of transfection. Fugene6 and Opti-MEM were warmed to room temperature and 6 μl of Fugene6 was diluted into 100 μl Opti-MEM, vortexed, and incubated 5 min at room temperature (RT). A total of 1.5 μg of plasmid DNA was added to the diluted Fugene6 at a 1:4 ratio (1.5 μg DNA/6 μl Fugene6), vortexed, and incubated for 15 min at RT. The transfection solution was then added drop-wise to cells and images were acquired 48 h post-transfection.

Blebbistatin, fasudil, H-1152, and ML7 were purchased from Tocris. Acrylamide and N,N^I^-methylenebisacrylamide were purchased from National Diagnostics. Tetramethyl-ethylenediamine (TEMED), ammonium persulfate (APS), and bovine serum albumin (BSA) along with other sundry chemicals were purchased from Sigma (St. Louis, MO). Plasmids used in this work include: PKA biosensor pmAKAR3 (43) from Dr. Jin Zhang (Johns Hopkins University); pYFP-dSH2 from Dr. Benny Geiger (Weizmann Institute of Science); pmCherry-FAK and pRFP-zyxin from Addgene (plasmid #35039 and 26720, respectively). The plasmid encoding mCherry-paxillin was made by substituting mCherry for EGFP in pEGFP-N1-paxillin (a gift from Dr. Chris Turner (SUNY, Upstate)), while the PKA inhibitor peptide fused to mCherry (mCherry-PKI) was described previously (32).

### Fabrication of polyacrylamide hydrogels

Acrylamide hydrogels with a Young’s elastic modulus of ~25 kPa were fabricated essentially as described previously (27). Briefly, cleaned 25 mm diameter round glass coverslips were briefly flamed and incubated with 0.1 N NaOH for 15 min. After removal of excess NaOH, 25 μl (3-aminopropyl)-trimethoxysilane (APTMS) was smeared on the coverslips and incubated for 3 min at RT and the coverslips were washed 3×5 min in ddH_2_O and dried by aspiration. Once dried, the coverslips were incubated with 500 μl 0.5% glutaraldehyde for 30 min. The glutaraldehyde was removed and a 25 μl drop of acrylamide solution (7.5:0.5% acrylamide:bis-acrylamide, activated with APS and TEMED), was sandwiched between the activated coverslip and a 22 mm diameter coverslip passivated with RainX (ITW Global brands, Houston, TX) and allowed to polymerize for 10 min. Once polymerized, the RainX-treated coverslip was removed and the hydrogel was washed three times (5 min each) in PBS. The gel surface was derivitized with the heterobifunctional cross-linker sulfo-SANPAH as previously described (27, 69). For routine studies, activated gels were functionalized with 20 μg/ml fibronectin at 37°C for 45 min. For traction force microscopy studies, 0.2 μm red-fluorescent carboxy-modified latex microspheres (Invitrogen, F8810) were conjugated to the gel surface by incubating a sonicated suspension of the beads (1:200 in 50 mM HEPES pH 8.5) on the gels for 30 min. The gels were rinsed 3x with 50 mM HEPES (pH 8.5) to remove all nonattached beads and then incubated with 20 μg/ml fibronectin (diluted in 50 mM HEPES pH 8.5) at 37°C for 45 min. The gels were post-fixed with 0.5% glutaraldehyde for 1 h at RT and quenched in NaBH4 before plating cells in complete media. Coated gels were washed 3 × 5 min in PBS and either used immediately or stored at 4°C for up to 1 week.

### Live cell imaging

Cells transfected with pmAKAR3 were cultured overnight in serum-free DMEM + 40 ng/ml epidermal growth factor (EGF), trypsinized, soybean trypsin inhibitor added, pelleted, and resuspended in DMEM 1% BSA + 40 ng/ml EGF. These cells were plated on fibronectin-coated coverslips and incubated for ~3 h prior to imaging at low density to induce migration. Cells plated on hydrogels for durotaxis were incubated overnight in complete media then rinsed twice in modified Ringer’s buffer without phosphate (10 mM HEPES; 10 mM Glucose; 155 mM NaCl; 5 mM KCl; 2 mM CaCl_2_; 1 mM MgCl_2_). All cells were re-fed modified Ringer’s buffer supplemented with 40 ng/ml EGF for imaging. Coverslips were mounted in a chamber (Attofluor, ThermoFisher) prior to imaging.

### Durotaxis assay

Cells were seeded on fibronectin-coated gels and mounted and maintained on the microscope as above. Cells were manipulated with a glass microneedle as described previously (32, 70). Briefly, micropipettes were fashioned from borosilicate glass capillaries (1B150–4 or TW150-4, World Precision Instruments) on a two-stage pipette puller (Pul-2, World Precision Instruments). A Narishige MF-900 microforge was used to form the micropipette tip into a hooked probe with a rounded end to engage the polyacrylamide hydrogels without tearing. The probe was mounted on a micro-manipulator (Leitz or Narishige) and lowered onto the gel surface approximately 20 μm away from a cell and pulled 20 μm in a direction orthogonal to the cell’s long axis. Quantification of response to stretch was calculated using custom ImageJ Protrusion-Retraction Analysis Mapping (PRAM) macros designed to calculate the percent of cell area protruding, retracting, and overlapping between any two given frames of time-lapse images (39). A positive durotactic response was defined as > 50% increase in protrusion index (protrusion area/(protrusion area + overlap area)) in the direction of stretch over 45 min, and durotactic efficiency was defined as the number of cells showing a durotactic response divided by the number of cells pulled.

### FRET Imaging and analysis

The FRET-based PKA activity biosensor, pmAKAR3, was imaged in SKOV-3 cells as previously described (32). Briefly, 48 h after transfection, cells were rinsed twice and maintained in HEPES-buffered saline solution containing (in mM): 134 NaCl, 5.4 KCl, 1.0 MgSO_4_, 1.8 CaCl_2_, 20 HEPES, and 5 D-glucose (pH 7.4) without serum, unless otherwise specified. Cells were imaged on a Nikon Eclipse TE-2000E inverted microscope with a 60×/1.4NA Plan Apo oil-immersion objective lens using the appropriate fluorophore-specific filters (Chroma Technology Corp, Rockingham, VT) and an Andor Clara charged coupled device camera (Andor Technologies, South Windsor, CT) controlled by Elements (Nikon) software. CFP, YFP and FRET images were acquired with 400–700 ms exposures and 2×2 binning for each acquisition at 60 sec intervals unless otherwise noted in the figure legends. Images in each channel were subjected to background subtraction, and FRET ratios were calculated using the BiosensorsFRET ImageJ plugin (71). Pseudocolor images were generated using a custom written ImageJ look-up table (72).

Leading edge PKA indices were generated by obtaining linescans form the cell centroid to the cell periphery in the direction of motility. The index is calculated from ~10 linescans from multiple cells by dividing the average FRET ratio of within 10 μm of the leading edge by the average FRET ratio within 10 μm of the cell centroid. This measurement both normalizes the FRET ratios so they can be compared between different cells and gives an accurate read-out of any change in PKA activity as a function of distance from the cell centroid.

### Cell Spreading and Migration assays

To monitor cell spreading, cells were prepared and cultured as previously described (42). Briefly, cells were serum starved overnight, trypsinized, quenched with 1mg/ml soybean trypsin inhibitor, washed via centrifugation (50 x g for 5 min), re-suspended in DMEM 1%BSA, and rocked for 1 h before plating on fibronectin coated (10 μg/ml) glass-bottom imaging dishes. The cells were allowed to settle to the bottom of the dish for 10 min at 4°C before imaging as described below. Similar conditions were used to monitor migrating cells with the exception that the cells were allowed to adhere and spread for 3 h at 37°C before imaging.

### Correlating edge velocity and PKA activity

Corrected FRET ratio time-lapse movies were fed to the Quantitative Imaging of Membrane Proteins (QuimP11) package (http://go.warwick.ac.uk/bretschneider/quimp) software that analyzed edge dynamics and calculated edge velocity. Additionally, the software generated two-dimensional morphodynamic plots of edge velocities along the cell edge over time and computed autocorrelation coefficients of edge dynamics. Once edge dynamics were analyzed, the corrected FRET ratio images were analyzed by QuimP11 to sample the FRET ratios within 10 μm from the cell edge. The software generated two-dimensional heat-maps of PKA activity along the cell edge over time and calculated cross-correlation coefficients at different time-lags to determine when peak PKA activity events were occurring in relation to peak protrusion events. The ImageJ plugin, QuimP11, was also used to display time coded depth stacks to depict cell movement over time in a single image.

### Focal adhesion analysis

Cells were transfected with plasmids encoding focal adhesion markers (mCherry-paxillin, mCherry-FAK, or RFP-zyxin) and either pmAKAR3 or dSH2-YFP to visualize focal adhesions and PKA activity or tyrosine phosphorylation. Images were acquired at 60 sec intervals. Focal adhesion pixel intensities and lifetimes were calculated by manually thresholding adhesions and measuring pixel intensities, adhesion assembly and disassembly rates, and lifetimes of individual adhesions in time-lapse movies. To quantify the spatiotemporal correlation between focal adhesion dynamics and PKA activity, leading edge focal adhesions were tracked over time and an interrogation region of interest (ROI) that was 50% larger than the area of the adhesion was used to track pmAKAR3 FRET ratios under the same cellular regions. Pearson’s correlation coefficients were generated using the Intensity Correlation Analysis ImageJ plugin, and average values are represented as mean ± SEM.

### Traction Force Microscopy

TFM was performed essentially as described previously (27). Briefly, cells were plated on polyacrylamide gels that were surface-conjugated with 0.2 μm red fluorescent latex microspheres as described above. Cells were adhered to the hydrogels overnight in complete media and were washed twice and maintained in HEPES-buffered saline as described above. Coverslips were mounted in imaging chambers as above and fluorescent beads images were captured though a 20x Plan Apo objective on a Nikon Eclipse TE-2000E inverted microscope as described above. Bead images were acquired before and after clearing cells by the addition of trypsin/EDTA (0.5%). Cell outlines were generated by either using the YFP image from cells expressing pmAKAR3 or a transmitted light image, in cases where cells were not transfected. Bead images were registered to correct for any stage drift, then the movement of individual microspheres between image pairs was calculated using Particle Image Velocimetry (PIV) and the Young’s elastic modulus of the polyacrylamide hydrogels (25 kPa) was used to calculate traction forces using Fourier Transform Traction Cytometry (FTTC; (73, 74)). The mean traction force within the cell was used to generate average cellular traction and the mean of the maximum traction forces was used to generate average maximum traction force generation.

### TFM/FRET correlation analysis

To correlate cellular traction forces and PKA activity, both readings were captured simultaneously and analyzed independently. Once heat maps of both PKA activity and traction forces were generated, standard image correlation analysis was performed. This analysis was made possible because the two signals are rendered as 8-bit gray-scale images and higher pixel intensity corresponds with either higher PKA activity or higher traction forces. Lookup tables are assigned to the images after analysis for ease of interpreting biosensor and TFM data. Mander’s correlation coefficients and intensity correlation quotients were generated using the Intensity Correlation Analysis ImageJ plugin, and average values are represented as mean ± SEM. Intensity correlation quotient (ICQ) analysis has been described in detail elsewhere. In brief, ICQ reflects the ratio of the number of positive (*Ai-aI*)(*Bi-b*) values to the total number of pixels in the region of interest, where *a* and *b* are the means of each signal intensity values *Ai* and *Bi*. ICQ values from - 0.05-+0.05 indicate random non-covariance between two signals, 0.05-0.1 indicate weak covariance, and >0.1 indicate strong covariance.

## Acknowledgements

We thank Drs. Benny Geiger (Weitzmann Institute), Jin Zhang (Johns Hopkins University), and Chris Turner (SUNY – Upstate) for generously providing plasmids. This work was supported by National Institute of General Medical Sciences (NIGMS) grants R01GM097495 and R01GM117490 (to A.K. Howe), NIGMS grant NIH R01GM097495-S1 (A.J. McKenzie), and a University of Vermont College of Medicine Pilot Project Award (A.K. Howe).

## Abbreviations list

AKAP: A-kinase anchoring protein
Blebb: Blebbistatin
*E*: Young’s elastic modulus
EOC: epithelial ovarian cancer
FRET: Förster resonance energy transfer
ICQ: intensity correlation quotient
PKA: cAMP-dependent protein kinase
mCh-PKI: mCherry-protein kinase inhibitor
pmAKAR3: plasma membrane A-kinase activity reporter 3
Pxn: paxillin
TFM: Traction force microscopy

## Online supplemental material

**Supplemental Figure S1.**
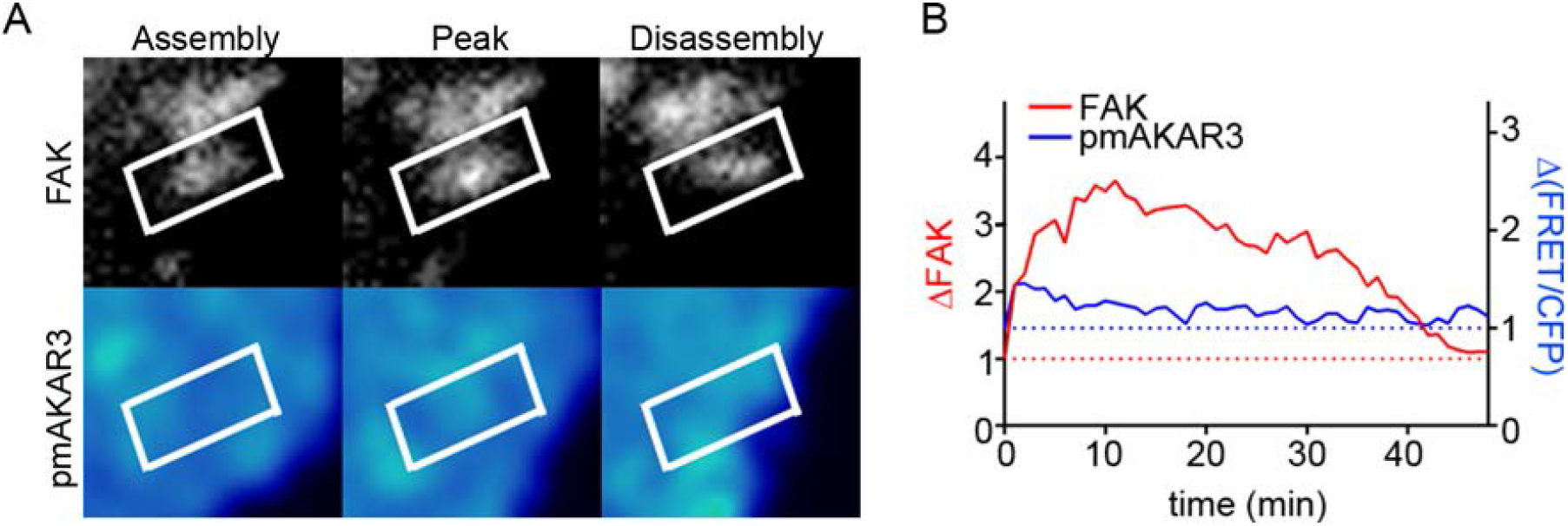
Lack of correlation between assembly of FAK-containing focal adhesions and localized PKA activity. (A) Migrating SKOV-3 cells co-expressing pmAKAR3 and mCherry-FAK were plated and imaged as described for Fig. 2. on fibronectin coated imaging dishes and representative near-simultaneous paxillin (Pxn) and pseudocolored FRET/CFP (pmAKAR3) images from live-cell microscopy experiments are shown. Images (15 min apart) of a single FAK-containing focal adhesion during assembly, peak, and disassembly (top) with overlapping pseudocolored FRET/CFP (bottom) are shown. (B) The changes in either FAK fluorescence intensity or FRET ratio in the indicated ROI were plotted over time.

**Supplemental Figure S2.**
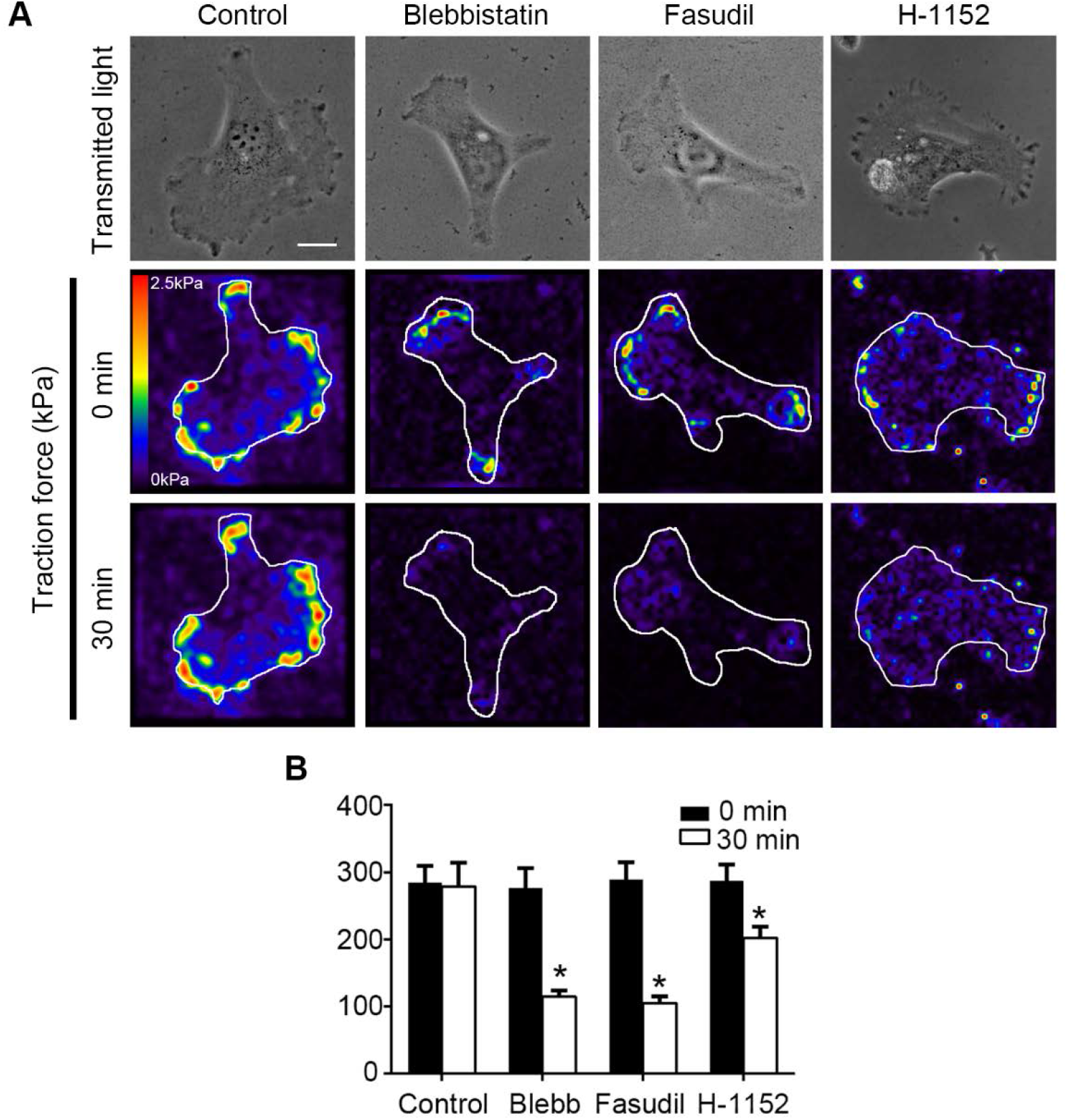
Cellular traction forces are inhibited with actomyosin contractility inhibitors and are low during cell spreading. (A) Transmitted light images and corresponding normalized traction force maps of cells before and 30 min after control, 25 μM Blebbistatin, 10 μM fasudil, or 1 μM H-1152 treatment. (B) Quantification of the average traction forces generated following indicated treatment (*n* = 10 cells for each condition; * = p< 0.01).

**Supplemental Figure S3.**
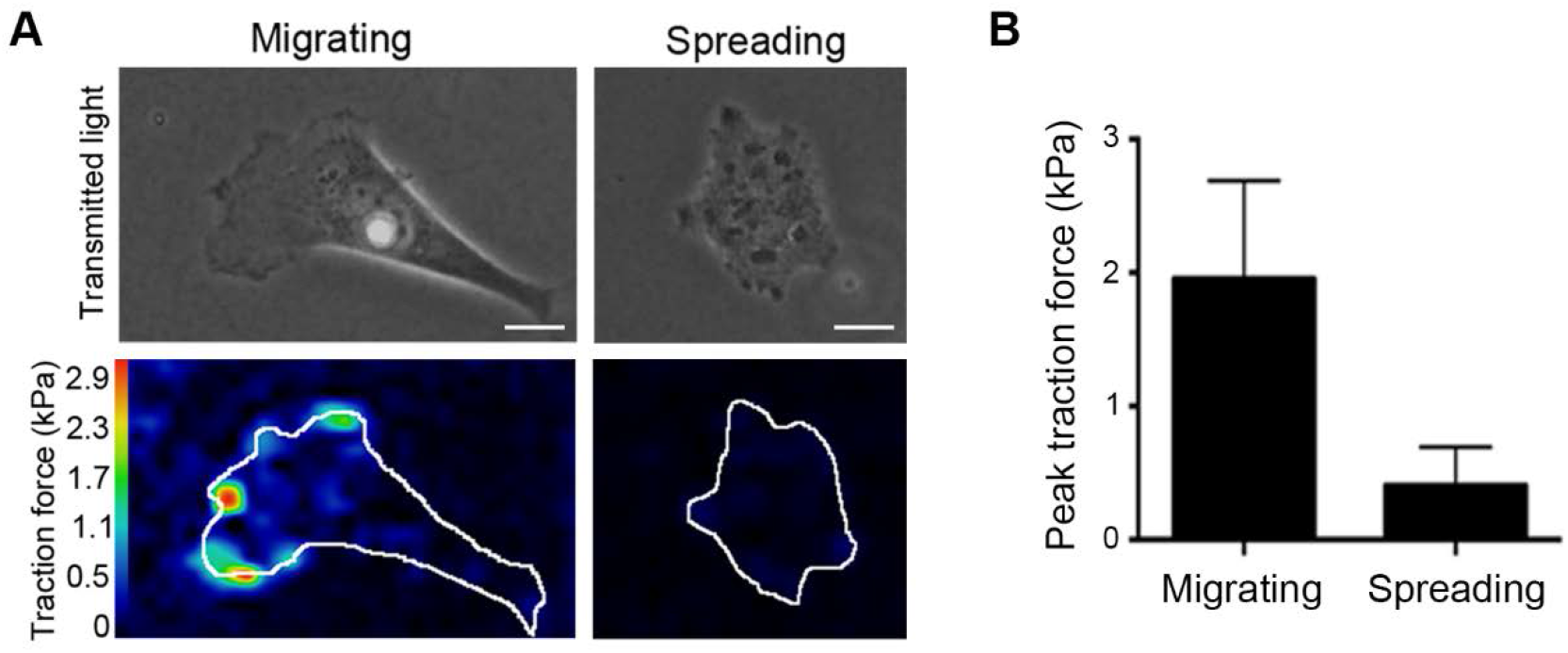
Spreading cells exhibit lower traction forces than migrating cells. (A) Transmitted light and normalized traction force maps of either migrating (left) or spreading (right) cells (bar = 10 μm). (B) Quantification of the average traction forces generated following indicated treatment (*n =* 4 for migration and *n* = 6 for spreading; * = p<0.01).

**Supplemental Movie S1. PKA is activated at the leading edge during cell migration but not at the periphery during cell spreading.** SKOV-3 cells expressing pmAKAR3 were plated on fibronectin-coated glass coverslips and monitored *via* live-cell FRET microscopy 10 min (*left*) or 4 h (*right*) after plating. Images were acquired every 60 sec for 12 min.

**Supplemental Movie S2. PKA is activated in the leading edge of SKOV-3 cells under a variety of migratory culture conditions.** Migrating SKOV-3 cells expressing pmAKAR3 were plated on fibronectin-coated glass coverslips and monitored *via* live-cell FRET microscopy under the following culture conditions: (a) serum-free medium overnight then serum-free media for imaging; (b) complete (serum-containing) medium overnight then serum-free media for imaging; (c) serum-free medium overnight then serum-free media with 40 ng/ml EGF added immediately before imaging; (d) serum-free medium overnight then serum-free media with 40 ng/ml EGF added 2 h before imaging. Images were acquired every 30 sec.

**Supplemental Movie S3. Leading edge PKA activity decreases upon treatment with blebbistatin.** An SKOV-3 cell expressing pmAKAR3 was imaged *via* FRET microscopy before and after treatment with 25 μM blebbistatin as indicated in the movie. Images were acquired every 2 min.

**Supplemental Movie S4. Leading edge PKA activity decreases upon treatment with Fasudil or ML-7.** SKOV-3 cells expressing pmAKAR3 were imaged *via* FRET microscopy before and after treatment with 10 μM fasudil (*left*) or 15 μM ML-7, as indicated in the movie. Images were acquired every 60 sec.

**Supplemental Movie S5. Rapid loss of zyxin from focal adhesions upon treatment with blebbistatin.** SKOV-3 cells expressing RFP-zyxin were imaged before and after treatment with DMSO (0.1% vol/vol) or 25 μM blebbistatin, as indicated in the movie. Images were acquired every 60 sec.

**Supplemental Movie S6. Rapid decrease in cellular traction force upon treatment with blebbistatin.** SKOV-3 cells were cultured on 125 kPa polyacrylamide hydrogels functionalized with fluorescent nanospheres. Bead fields were imaged before and after treatment with DMSO (0.1% vol/vol) or 25 μM blebbistatin, as indicated in the movie, and used to calculate cellular traction forces. Images were acquired every 60 sec.

**Supplemental Movie S7. Inhibition of PKA blocks durotaxis in SKOV-3 cells.** Control SKOV-3 cells or SKOV-3 cells expressing mCherry-PKI were plated on fibronectin-coated 25 kPa hydrogels and stretched with a glass micro-needle in the direction indicated by the white arrow. Images were acquired every 60 sec.

## References

1. Bershadsky AD, Balaban NQ, & Geiger B (2003) Adhesion-dependent cell mechanosensitivity. Annu Rev Cell Dev Biol 19:677–695.

2. Schiller HB & Fassler R (2013) Mechanosensitivity and compositional dynamics of cell-matrix adhesions. EMBO Rep 14(6):509–519.

3. Janmey PA & McCulloch CA (2007) Cell mechanics: integrating cell responses to mechanical stimuli. Annu Rev Biomed Eng 9:1–34.

4. Levayer R & Lecuit T (2012) Biomechanical regulation of contractility: spatial control and dynamics. Trends Cell Biol 22(2):61–81.

5. Ringer P, Colo G, Fassler R, & Grashoff C (2017) Sensing the mechano-chemical properties of the extracellular matrix. Matrix Biol 64:6–16.

6. Weinberg SH, Mair DB, & Lemmon CA (2017) Mechanotransduction Dynamics at the Cell-Matrix Interface. Biophys J 112(9):1962–1974.

7. Schwartz MA (2010) Integrins and extracellular matrix in mechanotransduction. Cold Spring Harb Perspect Biol 2(12):a005066.

8. Choquet D, Felsenfeld DP, & Sheetz MP (1997) Extracellular matrix rigidity causes strengthening of integrin-cytoskeleton linkages. Cell 88(1):39–48.

9. Balaban NQ, et al. (2001) Force and focal adhesion assembly: a close relationship studied using elastic micropatterned substrates. Nat Cell Biol 3(5):466–472.

10. Chrzanowska-Wodnicka M & Burridge K (1996) Rho-stimulated contractility drives the formation of stress fibers and focal adhesions. J Cell Biol 133(6): 1403–1415.

11. Lintz M, Munoz A, & Reinhart-King CA (2017) The Mechanics of Single Cell and Collective Migration of Tumor Cells. J Biomech Eng 139(2).

12. Parker KK, et al. (2002) Directional control of lamellipodia extension by constraining cell shape and orienting cell tractional forces. FASEB J 16(10): 1195–1204.

13. Plotnikov SV, Pasapera AM, Sabass B, & Waterman CM (2012) Force fluctuations within focal adhesions mediate ECM-rigidity sensing to guide directed cell migration. Cell 151(7):1513–1527.

14. Prager-Khoutorsky M, et al. (2011) Fibroblast polarization is a matrix-rigidity-dependent process controlled by focal adhesion mechanosensing. Nat Cell Biol 13(12):1457–1465.

15. Riveline D, et al. (2001) Focal contacts as mechanosensors: externally applied local mechanical force induces growth of focal contacts by an mDia1-dependent and ROCK-independent mechanism. J Cell Biol 153(6):1175–1186.

16. Roca-Cusachs P, Sunyer R, & Trepat X (2013) Mechanical guidance of cell migration: lessons from chemotaxis. Curr Opin Cell Biol 25(5):543–549.

17. Wolfenson H, Bershadsky A, Henis YI, & Geiger B (2011) Actomyosin-generated tension controls the molecular kinetics of focal adhesions. J Cell Sci 124(Pt 9):1425–1432.

18. Plotnikov SV & Waterman CM (2013) Guiding cell migration by tugging. Curr Opin Cell Biol 25(5):619–626.

19. Ingber DE (1997) Integrins, tensegrity, and mechanotransduction. Gravit Space Biol Bull 10(2):49–55.

20. Schiffhauer ES & Robinson DN (2017) Mechanochemical Signaling Directs Cell-Shape Change. Biophys J 112(2):207–214.

21. Pasapera AM, et al. (2015) Rac1-dependent phosphorylation and focal adhesion recruitment of myosin IIA regulates migration and mechanosensing. Curr Biol 25(2):175–186.

22. Vicente-Manzanares M, Newell-Litwa K, Bachir AI, Whitmore LA, & Horwitz AR (2011) Myosin IIA/IIB restrict adhesive and protrusive signaling to generate front-back polarity in migrating cells. J Cell Biol 193(2):381–396.

23. Beningo KA, Dembo M, Kaverina I, Small JV, & Wang YL (2001) Nascent focal adhesions are responsible for the generation of strong propulsive forces in migrating fibroblasts. J Cell Biol 153(4):881–888.

24. Bereiter-Hahn J & Luers H (1998) Subcellular tension fields and mechanical resistance of the lamella front related to the direction of locomotion. Cell Biochem Biophys 29(3):243–262.

25. Dembo M & Wang YL (1999) Stresses at the cell-to-substrate interface during locomotion of fibroblasts. Biophys J 76(4):2307–2316.

26. Lo CM, Wang HB, Dembo M, & Wang YL (2000) Cell movement is guided by the rigidity of the substrate. Biophys J 79(1):144–152.

27. McKenzie AJ, et al. (2018) The mechanical microenvironment regulates ovarian cancer cell morphology, migration, and spheroid disaggregation. Sci Rep 8(1):7228.

28. Howe AK (2004) Regulation of actin-based cell migration by cAMP/PKA. Biochim Biophys Acta 1692(2–3):159–174.

29. Howe AK (2011) Cross-talk between calcium and protein kinase A in the regulation of cell migration. Curr Opin Cell Biol 23(5):554–561.

30. Howe AK, Baldor LC, & Hogan BP (2005) Spatial regulation of the cAMP-dependent protein kinase during chemotactic cell migration. Proc Natl Acad Sci U S A 102(40): 14320–14325.

31. Paulucci-Holthauzen AA, et al. (2009) Spatial distribution of protein kinase A activity during cell migration is mediated by A-kinase anchoring protein AKAP Lbc. J Biol Chem 284(9):5956–5967.

32. McKenzie AJ, Campbell SL, & Howe AK (2011) Protein kinase A activity and anchoring are required for ovarian cancer cell migration and invasion. PLoS One 6(10):e26552.

33. Tkachenko E, et al. (2011) Protein kinase A governs a RhoA-RhoGDI protrusion retraction pacemaker in migrating cells. Nat Cell Biol 13(6):660–667.

34. Lim CJ, et al. (2007) Alpha4 integrins are type I cAMP-dependent protein kinase anchoring proteins. Nat Cell Biol 9(4):415–421.

35. Lim CJ, et al. (2008) Integrin-mediated protein kinase A activation at the leading edge of migrating cells. Mol Biol Cell 19(11):4930–4941.

36. Sinha C, et al. (2015) PKA and actin play critical roles as downstream effectors in MRP4-mediated regulation of fibroblast migration. Cell Signal 27(7):1345–1355.

37. Takahashi M, et al. (2013) Protein kinase A-dependent phosphorylation of Rap1 regulates its membrane localization and cell migration. J Biol Chem 288(39):27712–27723.

38. Zhang D, et al. (2010) Promotion of PDGF-induced endothelial cell migration by phosphorylated VASP depends on PKA anchoring via AKAP. Mol Cell Biochem 335(1–2):1–11.

39. Deming PB, et al. (2015) Anchoring of protein kinase A by ERM (ezrin-radixin-moesin) proteins is required for proper netrin signaling through DCC (deleted in colorectal cancer). J Biol Chem 290(9):5783–5796.

40. Giannone G, et al. (2007) Lamellipodial actin mechanically links myosin activity with adhesion-site formation. Cell 128(3):561–575.

41. Whittard JD & Akiyama SK (2001) Positive regulation of cell-cell and cell-substrate adhesion by protein kinase A. J Cell Sci 114(Pt 18):3265–3272.

42. Howe AK, Hogan BP, & Juliano RL (2002) Regulation of vasodilator-stimulated phosphoprotein phosphorylation and interaction with Abl by protein kinase A and cell adhesion. J Biol Chem 277(41):38121–38126.

43. Allen MD & Zhang J (2006) Subcellular dynamics of protein kinase A activity visualized by FRET-based reporters. Biochem Biophys Res Commun 348(2):716–721.

44. Mostafavi-Pour Z, et al. (2003) Integrin-specific signaling pathways controlling focal adhesion formation and cell migration. J Cell Biol 161(1):155–167.

45. Alenghat FJ, Tytell JD, Thodeti CK, Derrien A, & Ingber DE (2009) Mechanical control of cAMP signaling through integrins is mediated by the heterotrimeric Galphas protein. J Cell Biochem 106(4):529–538.

46. Goldmann WH (2002) The coupling of vinculin to the cytoskeleton is not essential for mechano-chemical signaling in F9 cells. Cell Biol Int 26(3):279–286.

47. O#Connor KL & Mercurio AM (2001) Protein kinase A regulates Rac and is required for the growth factor-stimulated migration of carcinoma cells. J Biol Chem 276(51):47895–47900.

48. Zaidel-Bar R, Ballestrem C, Kam Z, & Geiger B (2003) Early molecular events in the assembly of matrix adhesions at the leading edge of migrating cells. J Cell Sci 116(Pt 22):4605–4613.

49. Kirchner J, Kam Z, Tzur G, Bershadsky AD, & Geiger B (2003) Live-cell monitoring of tyrosine phosphorylation in focal adhesions following microtubule disruption. J Cell Sci 116(Pt 6):975–986.

50. Zhang X, et al. (2008) Talin depletion reveals independence of initial cell spreading from integrin activation and traction. Nat Cell Biol 10(9):1062–1068.

51. Wakatsuki T, Wysolmerski RB, & Elson EL (2003) Mechanics of cell spreading: role of myosin II. J Cell Sci 116(Pt 8):1617–1625.

52. Amano M, et al. (1996) Phosphorylation and activation of myosin by Rho-associated kinase (Rho-kinase). J Biol Chem 271(34):20246–20249.

53. Totsukawa G, et al. (2000) Distinct roles of ROCK (Rho-kinase) and MLCK in spatial regulation of MLC phosphorylation for assembly of stress fibers and focal adhesions in 3T3 fibroblasts. J Cell Biol 150(4):797–806.

54. Bosgraaf L & Van Haastert PJ (2010) Quimp3, an automated pseudopod-tracking algorithm. Cell Adh Migr 4(1):46–55.

55. Lavelin I, et al. (2013) Differential effect of actomyosin relaxation on the dynamic properties of focal adhesion proteins. PLoS One 8(9):e73549.

56. Lele TP, et al. (2006) Mechanical forces alter zyxin unbinding kinetics within focal adhesions of living cells. J Cell Physiol 207(1):187–194.

57. Hirata H, Tatsumi H, & Sokabe M (2008) Mechanical forces facilitate actin polymerization at focal adhesions in a zyxin-dependent manner. J Cell Sci 121(Pt 17):2795–2804.

58. Pasapera AM, Schneider IC, Rericha E, Schlaepfer DD, & Waterman CM (2010) Myosin II activity regulates vinculin recruitment to focal adhesions through FAK-mediated paxillin phosphorylation. J Cell Biol 188(6):877–890.

59. Li Q, et al. (2004) A syntaxin 1, Galpha(o), and N-type calcium channel complex at a presynaptic nerve terminal: analysis by quantitative immunocolocalization. J Neurosci 24(16):4070–4081.

60. Jaskolski F, Mulle C, & Manzoni OJ (2005) An automated method to quantify and visualize colocalized fluorescent signals. J Neurosci Methods 146(1):42–49.

61. Rodan GA, Bourret LA, Harvey A, & Mensi T (1975) Cyclic AMP and cyclic GMP: mediators of the mechanical effects on bone remodeling. Science 189(4201):467–469.

62. He Y & Grinnell F (1994) Stress relaxation of fibroblasts activates a cyclic AMP signaling pathway. J Cell Biol 126(2):457–464.

63. Meyer CJ, et al. (2000) Mechanical control of cyclic AMP signalling and gene transcription through integrins. Nat Cell Biol 2(9):666–668.

64. Funk SD, et al. (2010) Matrix-specific protein kinase A signaling regulates p21-activated kinase activation by flow in endothelial cells. Circ Res 106(8):1394–1403.

65. Hung WC, et al. (2016) Confinement Sensing and Signal Optimization via Piezo1/PKA and Myosin II Pathways. Cell Rep 15(7):1430–1441.

66. Goldfinger LE, et al. (2008) Localized alpha4 integrin phosphorylation directs shear stress-induced endothelial cell alignment. Circ Res 103(2):177–185.

67. Gui P, et al. (2006) Integrin receptor activation triggers converging regulation of Cav1.2 calcium channels by c-Src and protein kinase A pathways. J Biol Chem 281(20):14015–14025.

68. Storch U, Mederos y Schnitzler M, & Gudermann T (2012) G protein-mediated stretch reception. Am J Physiol Heart Circ Physiol 302(6):H1241–1249.

69. Tse JR & Engler AJ (2010) Preparation of hydrogel substrates with tunable mechanical properties. Curr Protoc Cell Biol Chapter 10:Unit 10 16.

70. Wang HB, Dembo M, Hanks SK, & Wang Y (2001) Focal adhesion kinase is involved in mechanosensing during fibroblast migration. Proc Natl Acad Sci U S A 98(20): 11295–11300.

71. Hodgson L, Shen F, & Hahn K (2010) Biosensors for characterizing the dynamics of rho family GTPases in living cells. Curr Protoc Cell Biol Chapter 14:Unit 14 11 11–26.

72. Stone RL, et al. (2014) Focal adhesion kinase: An alternative focus for anti-angiogenesis therapy in ovarian cancer. Cancer Biol Ther 15(7).

73. Tseng Q, et al. (2012) Spatial organization of the extracellular matrix regulates cell-cell junction positioning. Proc Natl Acad Sci U S A 109(5):1506–1511.

74. Marinkovic A, Mih JD, Park JA, Liu F, & Tschumperlin DJ (2012) Improved throughput traction microscopy reveals pivotal role for matrix stiffness in fibroblast contractility and TGF-beta responsiveness. Am J Physiol Lung Cell Mol Physiol 303(3):L169–180.

